# Neural field theory as a framework for modeling and understanding consciousness states in the brain

**DOI:** 10.1101/2024.10.27.619702

**Authors:** Daniel Polyakov, P.A. Robinson, Avigail Makbili, Olivia Gosseries, Oren Shriki

**Affiliations:** Department of Cognitive and Brain Sciences, Ben-Gurion University of the Negev, Be’er-Sheva, Israel; School of Physics, The University of Sydney, Sydney, NSW, Australia; Coma Science Group, GIGA Consciousness, University of Liège, Liège, Belgium; Centre du Cerveau, CHU of Liège, Liège, Belgium

## Abstract

Understanding the neural correlates of consciousness remains a central challenge in neuroscience. In this study, we explore the potential of neural field theory (NFT) as a computational framework for representing consciousness states. While prior research has validated NFT’s capacity to differentiate between normal and pathological states of consciousness, the relationship of its parameters to the representation of consciousness levels remains unclear.

Here, we fitted a corticothalamic NFT model to EEG data collected from healthy individuals and patients with disorders of consciousness. We then comprehensively explored the correlations between the fitted NFT parameters and features extracted from both experimental and simulated EEG data, across various states of consciousness. The identified correlations not only highlight the model’s ability to differentiate between states of consciousness, but also shed light on the physiological bases of these states, pinpointing potential biomarkers.

Our results provide valuable insights into how consciousness levels are represented within the NFT framework and into the dynamics of brain activity across various consciousness states. This underscores the potential of NFT as a useful tool for consciousness research, facilitating *in-silico* experimentation.

## Introduction

### Neural correlates of consciousness

Consciousness is a fascinating, hardly explained phenomenon. It exhibits a remarkable fluidity, transitioning in and out of our awareness throughout the day. We experience its return upon waking in the morning, as well as when we become drowsy after a satisfying meal. At times, we find ourselves drifting into a dreamlike state during moments of daydreaming. Paradoxically, it can entirely dissipate during deep sleep and then resurface during the enigmatic realm of dreams. Moreover, consciousness is subject to the influence of various substances, such as anesthetics and psychedelics, which can either augment or diminish its presence. Furthermore, it can be substantially impaired by an array of brain lesions and diseases, giving rise to disorders of consciousness (DoC), like coma or unresponsive wakefulness syndrome (UWS).

The study of consciousness has long captivated the imagination of philosophers, psychologists, and neuroscientists alike, igniting a profound quest to unravel the enigmatic neural underpinnings of this elusive phenomenon. This quest has given rise to an intriguing and multifaceted field of inquiry known as the “Neural Correlates of Consciousness” (NCC). The formation of the NCC field dates back approximately 30 years to the collaborative efforts of Nobel laureate Francis Crick and Christof Koch [1], and it has since experienced tremendous growth. NCC seek to elucidate the intricate relationship between subjective experience and the activity of the brain, striving to identify the specific neural mechanisms that give rise to our awareness of the world. This burgeoning field not only enhances our understanding of consciousness, but also holds the potential to shed light on a wide range of cognitive and clinical phenomena, ranging from transcendental meditation to absence epileptic seizures, ultimately deepening our insight into the fundamental nature of human existence.

### Consciousness-level quantification

Quantifying the level of consciousness poses a significant challenge within the domains of neuroscience and cognitive science. A plethora of methods and features have been developed to capture and measure this intricate aspect of human experience. Researchers often rely on behavioral observations and subjective self-report assessments to evaluate levels of consciousness, considering factors such as responsiveness, memory, and self-awareness. Another approach involves electrophysiological analysis of the brain through electroencephalography (EEG), intending to identify specific neural activity signatures associated with distinct states of consciousness.

EEG-based biomarkers of consciousness states typically rely on either power spectra or time-series complexity. Metrics like power in different frequency bands vary across wakefulness, sleep, anesthesia, and DoC. For instance, Delta and Theta band power increase with loss of consciousness, while higher Alpha and Gamma power are typical in wakeful states [2–4]. A more sophisticated metric is the spectral exponent *β*, or the spectral slope on a logarithmic scale, which is derived from the power law behavior of the EEG spectrum *P* (*f*) ∝ *f^−β^*, also termed “1*/f* spectrum”. Research indicates that this slope tends to be steeper during sleep and anesthesia compared to wakefulness [5–8]. Spectral entropy, which estimates the complexity of the power spectrum, also holds promise for estimating consciousness states, with higher values typically observed in less severe DoC states [2, 9]. For assessing time-series complexity, Lempel-Ziv complexity (LZc) is commonly employed, measuring the rate at which unique patterns appear in brain activity. LZc analysis of resting-state EEG has proven capable of distinguishing between anesthetic, psychedelic, and wakeful states, with higher conscious states exhibiting greater EEG complexity [8, 10–12]. Moreover, LZc analysis of EEG signals acquired after transcranial magnetic stimulation has successfully differentiated among DoC states [13]. Conversely, permutation entropy (PE) assesses the degree of disorder or randomness within the time series of neural signals. PE has been effectively utilized in epilepsy research and demonstrated the ability to discriminate between various sleep stages, with lower PE values corresponding to deeper sleep stages [14, 15].

### Disorders of consciousness

DoC encompass a range of conditions characterized by impaired awareness and wakefulness resulting from severe brain damage. These conditions include coma, UWS, and minimally conscious state (MCS) [16]. Within the MCS category, patients are often further classified as MCS- or MCS+ based on their nonreflex responses, verbal abilities, and command, following [17]. Additionally, the term “emergence from MCS” refers to the condition when MCS patients regain functional communication or object use [18], although this is not strictly classified as DoC. Collectively, these conditions disrupt the intricate neural processes that underlie human consciousness irrespective of arousal, challenging our fundamental understanding of how the brain forms awareness, and cognition, thereby prompting profound scientific, clinical, and ethical inquiries [16, 19].

In the present work, we analyze a dataset of patients with DoC (see the “Experimental data” section under Methods). This particular dataset is advantageous because it encompasses five distinct and clearly defined levels of consciousness. In contrast to binary conscious/unconscious datasets, a multi-level dataset offers the opportunity to test for more refined classification capabilities and provides a detailed interpretation of various DoC states. Additionally, it enables us to treat the consciousness level as an approximately continuous function, which is useful for conducting correlation analyses.

### Brain activity model for consciousness representation

Over the past half-century, substantial progress has been made in the development of brain activity models, addressing a wide range of phenomena occurring across different brain regions and scales [20–23]. A significant focus of these efforts has been the creation of comprehensive, physiology-based, multiscale models that can simulate various brain phenomena and produce realistic signals. Finding such a model, that is capable of reliably representing different levels and states of consciousness can be very helpful in many research aspects in this field. Such a model would be immensely valuable for interpreting various phenomena, generating hypotheses, and conducting *in-silico* experiments. For instance, consider the scenario of designing a novel invasive therapy for patients with DoC. First, we can fit the model to each individual patient with DoC, and the model’s parameters can be explored to gain a deeper understanding of the physiological condition of each patient. Subsequently, a personalized therapy plan can be proposed for each patient, based on the state of their fitted model. Following this, we can perform a simulation of the proposed therapy applied to the model, with the generated output signal serving as a tuning mechanism and a biomarker for the desired therapeutic effect [24]. Only after a successful outcome in the virtual model can these strategies be cautiously extended to real-life patient trials. This approach minimizes risks and optimizes the potential for therapeutic success in a highly controlled and personalized manner.

We have evaluated various models that hold the potential to fulfill this purpose. Among them, the most renowned is the “Blue Brain Project”, which seeks to construct a precise cortical network by linking detailed single-neuron models with anatomically derived connections. However, the project has encountered computational limitations, particularly when attempting to simulate large neuronal populations, let alone the entire brain [21]. A more pragmatic alternative is “The Virtual Brain,” which represents brain regions through neural mass models that are interconnected based on connectome data. This model has the capability to simulate diverse phenomena such as epilepsy and sleep stages. Nonetheless, the challenge lies in fitting it to experimental data due to the utilization of non-linear differential equations to generate the activity of each region [22, 25]. In contrast, neural field theory (NFT) modeling offers an efficient approach to constructing networks of interconnected brain regions that interact through wave equations. The NFT framework provides convenient tools for fitting and simulating neural activity using standard computing equipment [20, 26, 27]. Furthermore, NFT has shown efficacy in representing various consciousness-related phenomena, such as sleep stages and epilepsy [28–30], and it has also been applied to classify DoC states [31]. This makes NFT an attractive candidate for creating models that can deepen our understanding of consciousness and related disorders.

### Research motivation

In this study, we aim to assess the potential of NFT modeling as a viable tool in the research of NCC. Through our work, we not only replicate findings from prior studies, but also furnish evidence supporting theoretical predictions. Additionally, we unveil numerous new abilities and characteristics inherent to the NFT model. Our results indicate that the level of consciousness is reflected in specific model parameters and the signals produced by the model. In essence, our study demonstrates that NFT serves as a reliable model for NCC research, paving the way for future advancements in this field.

Going into detail, in our work, we exploit the ability of NFT to be easily fitted to experimental EEG data and to, subsequently, generate artificial EEG signals accordingly. We then extract a variety of features from both the experimental and simulated EEG data. This process is applied to a dataset of patients with DoC, where we seek to identify correlations between these features, the model parameters, and the consciousness states of the patients. Additionally, we show that fitted NFT parameters can be employed for the classification of consciousness states, as well as for their physiological interpretation. The parameters also offer insights into the interpretation of features based on simulated EEG that are correlated with them. The correlations observed between experimental-EEG-based and simulated-EEG-based features suggest that a fitted NFT model can reproduce and control them. The knowledge derived from these various correlations enables us to ascertain the capability of NFT to represent consciousness, reinforcing its value as a tool for investigating this fascinating aspect of the human brain.

## Materials and methods

### Neural field theory

Neural-field modeling [20, 32, 33] stands as a tool for constructing physiologically inspired brain models capable of mimicking various multiscale measures of brain activity. This approach captures a continuum of corticothalamic activity by simulating the local dynamics in each population and employing wave equations to describe the propagation between these populations. [34]. Consequently, it effectively replicates and integrates numerous phenomena observed in EEG data, including spectral peaks observed during waking and sleeping states, event-related potentials, measures of connectivity, and spatiotemporal structure, and even the dynamics of epileptic seizures (see Figure 1) [29, 35–41]. The model’s parameters encompass various biophysically meaningful quantities such as synaptic strengths, excitatory and inhibitory gains, propagation delays, synaptic and dendritic time constants, and axonal ranges. NFT represents a bottom-up approach to whole-brain modeling, which involves averaging over microstructure to derive mean-field equations. This method effectively complements analyses conducted at the cellular and local neural network levels. Additionally, NFT’s strength lies in its dynamic analysis, which explores the model’s states and their stability in relation to changes in model parameters [26, 39, 40].

**Figure 1.**
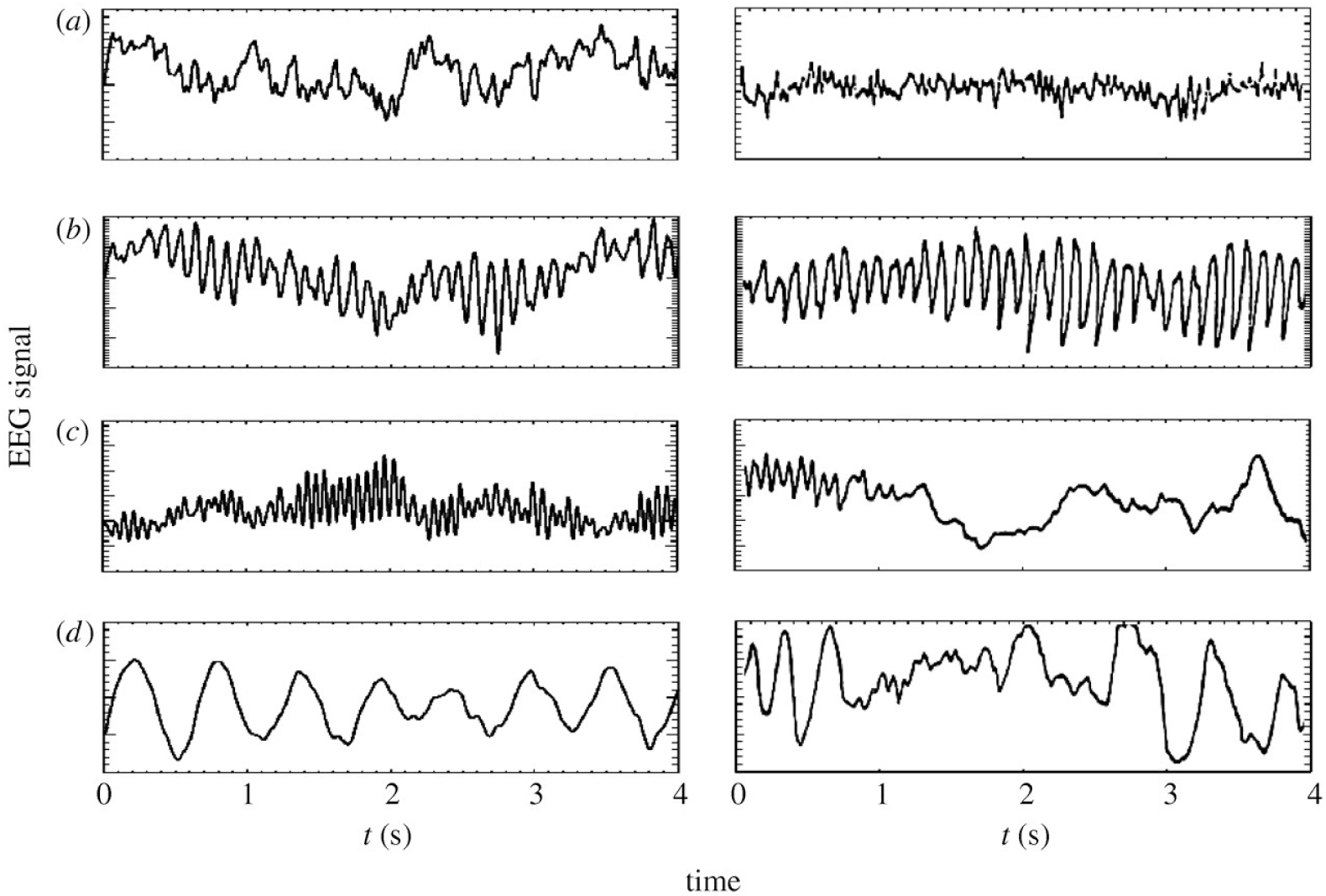
Simulated VS experimental EEG time series of waking and sleeping states. In the left panels: model generated time series of (a) eyes-open resting state, (b) eyes-closed resting state, (c) sleep-stage 2, and (d) sleep-stage 3. In the right panels: corresponding time series from human subjects [50, 51]. (The figure was originally published by Robinson et al. (2005) [20].)

The neural field *ϕ*(**r***, t*) [s*^−^*^1^] represents a spatiotemporal neural activity propagating among neural populations when averaged across scales of approximately 0.1 millimeters. Within the framework of the CTM, four distinct neural populations are involved, with key connectivities, illustrated schematically in Figure 2. These populations comprise excitatory (*e*) and inhibitory (*i*) cortical neurons, thalamic relay nuclei neurons (*s*), thalamic reticular nucleus neurons (*r*), and sensory inputs (*n*). Each of these populations has a soma potential *V_a_*(**r***, t*) [V] that is influenced by contributions *ϕ_b_* from presynaptic populations, and generates outgoing neural activity *ϕ_a_*(**r***, t*).

**Figure 2.**
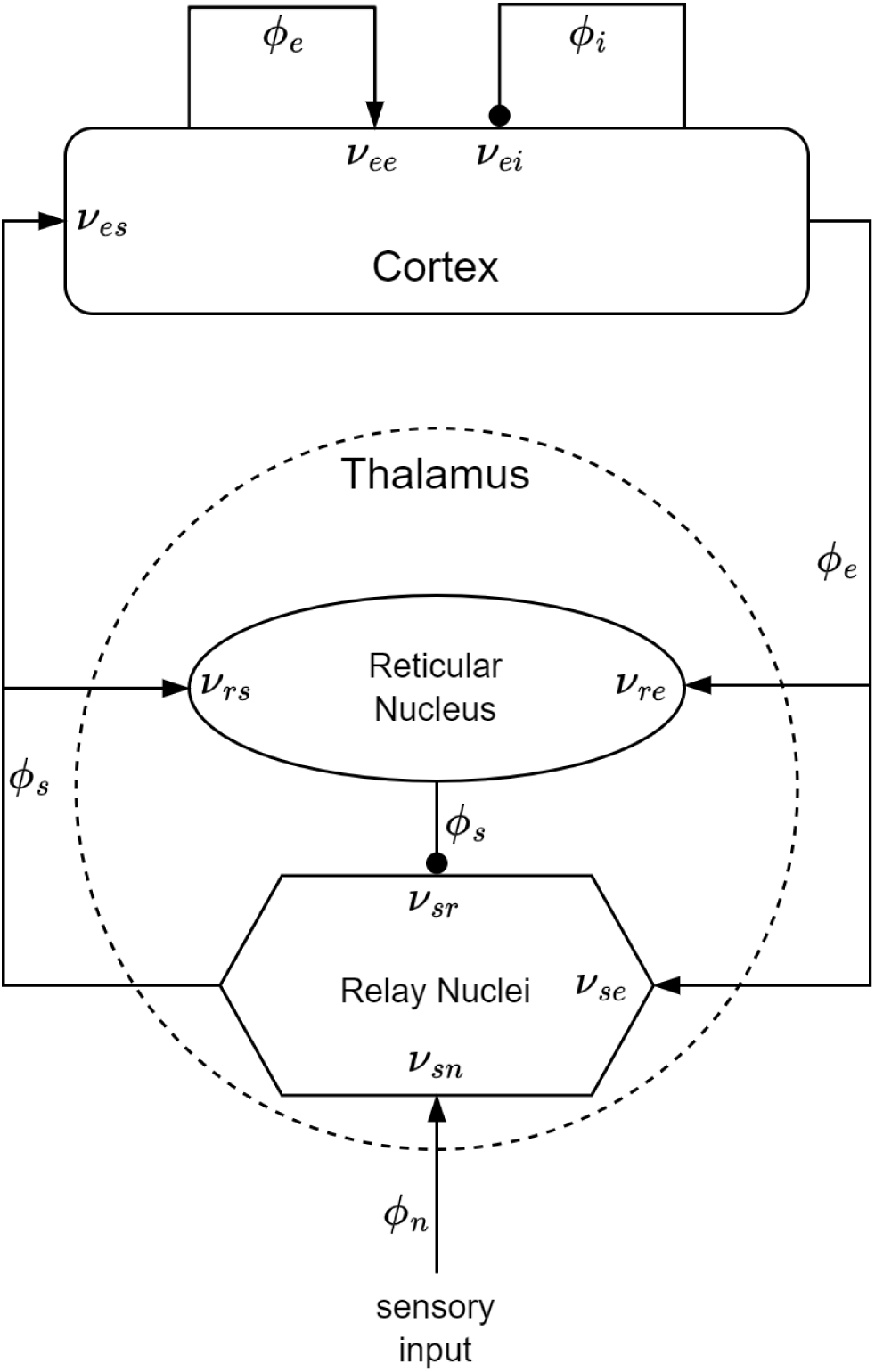
The CTM diagram. The neural populations shown are cortical excitatory, *e*, and inhibitory, *i*, thalamic reticular nucleus, *r*, and thalamic relay nuclei, *s*. The parameter *ν_ab_* quantifies the strength of the connection from population *b* to population *a*. Excitatory connections are indicated by pointed arrowheads, while inhibitory connections are denoted by round arrowheads.

The dendritic spatiotemporal potential *V_ab_*[V] is linked to the input *ϕ_b_* through Eq. 1. The parameter *ν_ab_* = *s_ab_N_ab_* [V · s] represents the strength of the connection from populations *b* to *a*, where *N_ab_* is the mean number of synapses per neuron *a* from neurons of type *b*, and *s_ab_* [V · s] is the mean time-integrated strength of soma response per incoming spike. The parameter *τ_ab_* [s] refers to the one-way corticothalamic time delay, and *D_a_*(*t*) is a differential operator, as described in Eq. 2. Here, 1*/α* and 1*/β* denote the characteristic decay time and rise time, respectively, of the soma response within the corticothalamic system.

Dendritic component:

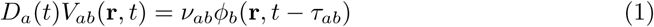

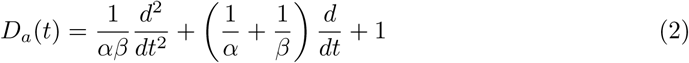

The soma potential *V_a_* is determined as the sum of its dendrite potentials, as outlined in Eq. 3. This potential undergoes some smoothing effects attributed to synaptodendritic dynamics and soma capacitance. Furthermore, the population generates spikes at a mean firing rate *Q_a_* [s*^−^*^1^], which is related to the soma potential through a sigmoid function *S*(*V_a_*) (relative to the resting state), as shown in Eq. 4. In this equation, *Q*_max_ denotes the maximum firing rate, while *θ* and 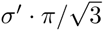 correspond to the mean and the standard deviation (SD), respectively, of the firing threshold voltage.

Soma component:

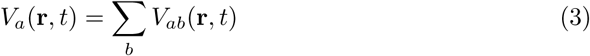

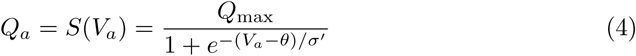

The field *ϕ_a_* approximately follows a damped wave equation with a source term *Q_a_*, as detailed in Eq. 5. The differential operator *D_a_*(**r***, t*) is defined in Eq. 6, where *v_a_* [m · s*^−^*^1^] represents the propagation velocity, *r_a_* [m] denotes the mean range, and *γ_a_* = *v_a_/r_a_* [s*^−^*^1^] signifies the damping rate.

Axonal component:

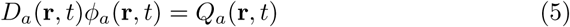

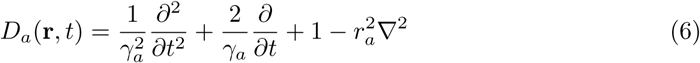

In this model, only *r_e_* is large enough to induce notable propagation effects. Consequently, the fields of other populations can be approximated as *ϕ_a_*(**r***, t*) = *S*[*V_a_*(**r***, t*)]. Additionally, we assume that the only non-zero time delays between populations are *τ_es_*, *τ_is_*, *τ_se_*, and *τ_re_*= *t*_0_*/*2, where *t_0_* is the total time it takes to traverse the corticothalamic loop. It is important to note that Eq. 5 encompasses the corticocortical time delays, as the wave equation inherently accounts for delays arising from propagation across the cortex. To further simplify the model, we assume random intracortical connectivity, leading to *N_ib_* = *N_eb_* for all *b* [42]. This assumption implies that the connection strengths are also symmetric, resulting in *ν_ee_* = *ν_ie_*, *ν_ei_* = *ν_ii_*, and *ν_es_* = *ν_is_* [20, 26]. Numerical integration [27] or, when feasible, analytical integration [43] of NFT equations produces a spatiotemporal activity signal that propagates across the cortical surface.

### CTM EEG power spectrum

In scenarios of spatially uniform steady-state activity, it is possible to analytically compute the power spectrum of the model, eliminating the necessity for numerical integration. The steady state is attained by setting all time and space derivatives to zero. Employing the first term of the Taylor expansion enables a linear approximation of all potential perturbations from the steady state. Applying a Fourier transform to the model equations under these conditions yields Eq. 7 for the dendritic component and Eq. 8 for the axonal component. Within these equations, *ω*= 2*πf* denotes the angular frequency, *k* = 2*π/λ* signifies the wave vector (*λ* is the wavelength), and *V* ^(0)^ represents the steady-state potential [26].

Dendritic equations in the frequency domain:

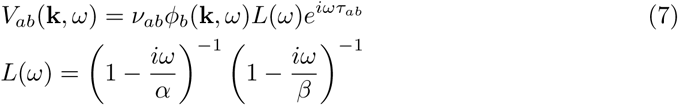

Axonal equations in the frequency domain:

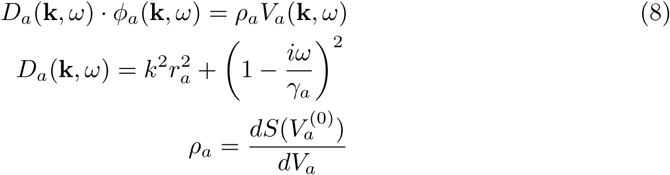

Using Eq. 3, we can write Eq. 8 as:

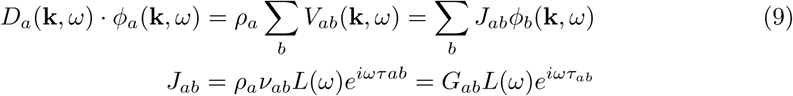

Then we can represent the interactions among the different populations within the CTM in matrix form:

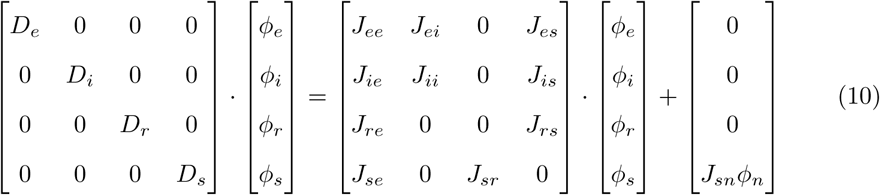

Eq. 10 can also be written in a compact form, when **J\**ϕ*\*** is the external input to the CTM:

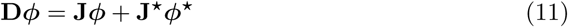

By solving Eq. 10, considering all the previously mentioned assumptions regarding *D_a_*, *ν_ab_*, and *τ_ab_*, we can derive Eq. 12. In this context, the quantities *G_ese_* = *G_es_G_se_*, *G_esre_* = *G_es_G_sr_G_re_*, and *G_srs_* = *G_sr_G_rs_*correspond to the overall gains for the excitatory corticothalamic, inhibitory corticothalamic, and intrathalamic loops, respectively. The firing rate of sensory inputs to the thalamus, *ϕ_n_*, is approximated by white noise. Without loss of generality, *ϕ_n_*(*ω*) can be set to 1, while only *G_sn_*is subject to variation.

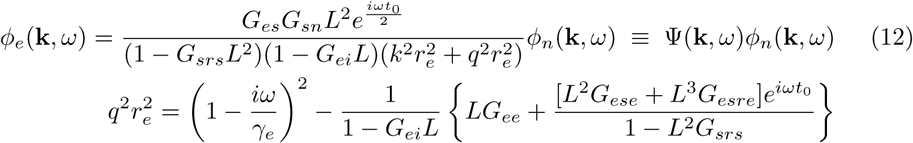

The excitatory field *ϕ_e_* is considered a good approximation of scalp EEG signals [26]. The EEG power spectrum *P* (*ω*) (Eq. 13) is calculated by integration of |*ϕ_e_*(**k***, ω*)|^2^ over **k** when the cortex is approximated as a rectangular sheet of size *L_x_* × *L_y_*. When considering periodic boundary conditions, this integral transitions into a summation over spatial modes with a discrete *k*. The filter function *F* (*k*) serves as an approximation of the low-pass spatial filtering that occurs due to volume conduction through the cerebrospinal fluid, skull, and scalp.

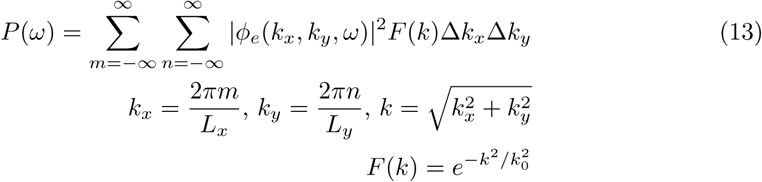

### CTM stability

Prior research has demonstrated that the CTM can effectively describe both healthy and pathological brain activity. Healthy brain activity, such as wake–sleep cycles, corresponds to stable modes of the model, while pathological conditions, like epileptic seizures, are associated with unstable modes [28, 29].

The stability of the CTM is determined by the denominator in Eq. 12. When all the zeros of Eq. 14 satisfy *Im*(*ω*_0_) *<* 0, the model is stable. Model stability boundaries can be represented in a three-dimensional space, as defined in Eq. 15. The *X,Y,Z* parameters correspond to corticocortical, corticothalamic, and intrathalamic loop strengths, respectively. These parameters offer a qualitative representation of the majority of CTM’s dynamics and provide insights into its stability characteristics across various parameter values [30].

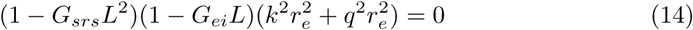

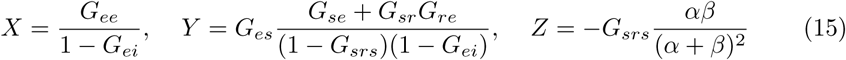

A notable stability boundary in the CTM is represented by the equation *X* + *Y* = 1. At this boundary, a saddle-node bifurcation occurs, and the model exhibits the generation of epileptic seizures [28]. Interestingly, prior research has also demonstrated that both *X* + *Y* and *X* − *Y* can be utilized to distinguish between different states of consciousness [31].

### Experimental data

In our study, we utilized a dataset consisting of open-eyes resting state EEG recordings acquired from a total of 117 DoC patients and 23 healthy control subjects. The etiology of DoC patients included traumatic brain injury (TBI), stroke, anoxia, cardiac arrest, and other causes. The severity of DoC was assessed using the Coma Recovery Scale-Revised (CRS-R) [18] several times within a few days, including just before the EEG acquisition. All patients were medically stable, breathing spontaneously, and resided in hospitals, rehabilitation centers, or at home. The DoC patient group comprised 28 UWS patients (12 females, ages 24–79, CRS-R 4–9, 6 had TBI, 12–173 months since injury), 20 MCS-patients (11 females, ages 21–80, CRS-R 7–13, 9 had TBI, 36–129 months since injury), 50 MCS+ patients (19 females, ages 11–78, CRS-R 8–20, 23 had TBI, 11–196 months since injury), and 19 patients in emergence from MCS (9 females, ages 17–73, CRS-R 16–23, 10 had TBI, 18–94 months since injury). The healthy control group consisted of 11 females and 12 males, aged between 19 and 69 years. Each participant’s EEG activity was recorded using a Hydro-Cel GSN high-density electrode net (Electric Geodesics, EGI) equipped with 256 electrodes, with a sampling rate of 250Hz. Data collection took place in a controlled environment, characterized by darkness and minimal sensory stimuli, with recording sessions lasting approximately 30 minutes [44].

### EEG processing

Each subject’s EEG data went through several processing steps. First, we kept only 137 central electrodes and removed the rest in order to focus only on scalp-based electrodes. The signals from these electrodes were high-pass filtered above 0.2Hz and notched at 50Hz to eliminate line noise. Subsequently, to streamline computational processes, they were down-sampled to 125Hz. The 30-minute recording was then partitioned into nearly stationary sections (required for CTM fitting, which is performed through steady-state activity power spectrum), with durations varying from 2.5 to 10 minutes. Detecting potential onsets of stationary sections involved splitting the electrode signals into 10-second segments and identifying when the segment’s SD deviated by two SDs compared to the preceding segment. Defining stationary section boundaries required simultaneous onsets in more than 5% of the electrodes, with at least 2.5 minutes between onsets. Sections lasting between 10 and 20 minutes were equally divided into two, while those between 20 and 30 minutes were split into three sections.

The cleaning process for each stationary section of the data involved several sequential steps. Initially, noisy channels (electrode signals) were identified by segmenting the data into 5-second segments and calculating the SD of each channel within a segment. A channel within a segment was flagged as noisy if its SD exceeded seven times the SD of the entire non-segmented channel, or if it was three times higher than that of the other channels within the same segment. Segments with at least one noisy channel were deemed noisy, and entire channels were marked as noisy if more than 33% of their segments exhibited noise. These identified entire noisy channels were then removed and replaced with channels generated through a spherical spatial interpolation of the remaining channels. Subsequently, the data was re-referenced to the average of all channels and any remaining artifacts were eliminated. Artifacts were defined as 1-second segments where any sample’s SD was five times greater than the channel’s entire SD. To further refine the data, we conducted an independent component analysis to isolate and remove components associated with eye movements and non-EEG activity. After decomposition, we employed the *IClabel* toolbox [45] to retain only those components identified with at least 50% confidence as originating from genuine neural activity. The steps involving spatial interpolation, average referencing, and independent component analysis were all executed using the *EEGLAB* toolbox [46].

From the various stationary sections of each subject, we retained sections where fewer than 10% of the channels were flagged as noisy and fewer than 10% of the independent components were removed during the cleaning process. Among these sections, we selected the one with the least amount of detected artifacts per minute of recording. Then, we calculated the power spectra of all EEG channels in this selected stationary section using a fast Fourier transform, to be further used for CTM fitting.

### Model fitting and signal generation

The CTM-fitting procedure was conducted by adjusting the analytical form of the CTM power spectrum (Eq. 13) to match the power spectra of the stationary section within a frequency range of 1–40Hz. This fitting process was carried out using a Monte-Carlo Markov-Chain optimization procedure applied to model parameters, a method implemented in the *braintrak* toolbox [30]. We optimized the following model parameters: *G_ee_*, *G_ei_*, *G_es_*, *G_se_*, *G_sr_*, *G_sn_*, *G_re_*, *G_rs_*, *α*, *β*, *t_0_*, and *EMG_a_*. To support multi-electrode fitting, we assumed that *G_ee_*, *G_ei_*, *G_sn_*, *α*, *β*, and *t_0_* exhibit a cosine-like variation over the scalp [34].

Time series data were generated from the fitted model by numerically integrating its partial differential equations over time, incorporating the *NFTsim* toolbox [27]. The resulting artificial EEG signals, with a duration of 2.5 minutes, were sampled at 125Hz and appeared at 784 locations on a 0.5 m × 0.5 m square sheet. These signals were then interpolated to match the 137 experimental electrode locations, high-pass filtered above 0.2Hz, and notched at 50Hz, as we did in experimental data processing.

An illustrative example showing an experimental EEG Cz electrode power spectrum, a power spectrum derived from the fitted model, and a simulated EEG Cz electrode power spectrum is depicted in Figure 3. The Cz electrode was specifically chosen to illustrate spectra of simulated data throughout this work, as its activity effectively represents the overall EEG activity generated by the CTM.

**Figure 3.**
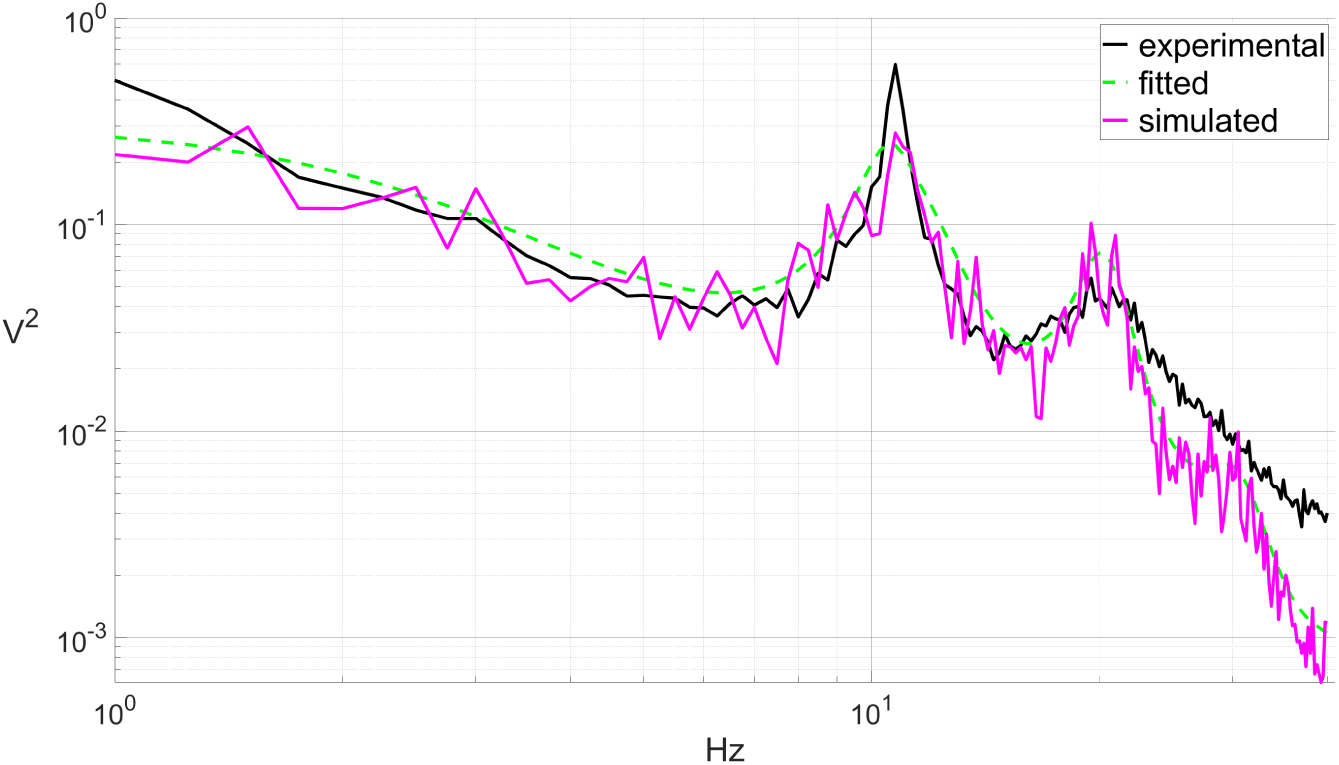
Experimental, fitted, and simulated power spectra. Example of a Cz electrode EEG power spectrum of a healthy subject, a fitted CTM power spectrum, and a Cz electrode power spectrum of a simulated EEG.

### Feature extraction

Feature extraction plays a crucial role in our study, serving multiple purposes. First, these features are employed to gauge the level of consciousness in the subjects and their corresponding (fitted) CTMs. Next, we investigate correlations between features extracted from the experimental EEG and those derived from simulated EEG to evaluate how well a fitted CTM reproduces them. Lastly, we explore the impact of NFT parameters on various features by analyzing their interrelations. For each subject, the following features were extracted from both the experimental and the simulated EEG:

#### Spectrum-related features

For each EEG channel, we computed a normalized power spectrum and calculated the total power within the typical EEG bands: Delta (1–4Hz), Theta (4–8Hz), Alpha (8–12Hz), Beta (12–24Hz), and Gamma (24–40Hz). Additionally, we calculated the spectral slope (the spectral exponent *β*) within the frequency range of 1 to 8 Hz using the FOOOF toolbox [47]. Furthermore, we computed the spectral entropy of each EEG channel using the algorithm introduced by Piarulli et al. (2016) [2]. Despite being an entropy-related feature, we mention it here as it is derived from the power spectrum.

#### Entropy-related features

We calculated the permutation entropy (PE) from the time series of each EEG channel using Ouyang’s toolbox [14]. Additionally, we extracted Lempel-Ziv complexity (LZc) from all channels collectively using Fekete’s toolbox [11].

While all spectrum-related features and PE were computed for each EEG electrode, we opted to showcase the results specifically for the Cz electrode throughout this study. The Cz electrode effectively captures the overall EEG activity simulated by the CTM. On the other hand, experimental EEG activity may vary across the scalp; however, this variability is captured by multi-electrode fitting. Therefore, we allow ourselves to use this electrode to present experiment-based features as well, sacrificing some accuracy but gaining consistency with simulation-based features.

### Parameter-feature correlation and sensitivity analysis

In order to understand how each NFT parameter influences the features extracted from simulated EEG, we performed a correlation analysis. This analysis involved calculating the Pearson correlation between different parameters and features, using data from all 140 subjects. However, it was necessary to ensure that the computed correlations were not a result of combinations of other NFT parameters. To address this, we also performed a comprehensive grid search across the parameter space and examined the sensitivity of features to various parameter values. To this end, we selected two representative subjects, one healthy and one MCS-, whose power spectra closely resembled the median power spectra of all subjects in the same state. We then utilized their fitted CTM to generate EEG signals and extract features while systematically altering the parameters one at a time. Each parameter was sampled from the entire possible parameter range while keeping the other parameters at their original fitted values. The altered parameters and their respective ranges are detailed in Table 1.

**Table 1.** Altered parameters and their ranges.

| **Parameter** | **Description** | **Range** |
| --- | --- | --- |
| $G_{ee}$ | Excitatory cortical gain | 0-20 |
| $G_{ei}$ | Inhibitory cortical gain | (-20)-0 |
| $G_{es}$ | Thalamic relay nuclei to cortex connection gain | 0-11 |
| $G_{se}$ | Cortex to thalamic relay nuclei connection gain | 0-17 |
| $G_{sr}$ | Reticular to relay intrathalamic nuclei connection gain | (-9)-0 |
| $G_{re}$ | Cortex to thalamic reticular nuclei connection gain | 0-7 |
| $G_{rs}$ | Relay to reticular intrathalamic nuclei connection gain | 0-8 |
| $\alpha$ | Inverse synaptodendritic decay time | 10-120 |
| $\beta$ | Inverse synaptodendritic rise time | 100-800 |
| $t_0$ | Corticothalamic loop delay | 0.075-0.14 |

We computed parameter-feature correlations based on the sensitivity analysis of the two representative subjects and then compared them to ones acquired during an all-subjects correlation analysis. The goal was to identify whether there were similar or opposite trends in each parameter-feature correlation. With that said, it is important to note that the examination of only two subjects is not sufficient. To ensure the robustness of parameter-feature correlations, this procedure should ideally be conducted across the entire dataset. In this study, we illustrate the feasibility of trends comparison based on a sensitivity analysis using these representative subjects, recognizing that a comprehensive inspection of trends for every subject in the dataset is a substantial undertaking that exceeds the scope of our current work.

## Results

We utilized NFT to model levels of consciousness. By fitting a CTM to EEG data from 23 healthy subjects and 117 DoC patients, we investigated the correlations between fitted NFT parameters and features extracted from experimental and simulated EEG. Our analysis revealed significant correlations between various features and NFT parameters, particularly noting strong associations with stability-related parameters (*X*, *Y*, etc.), shedding light on the physiological basis of NFT parameters. By matching healthy and various DoC states to these correlations, we have demonstrated the capabilities of NFT in representing states of consciousness.

### Discrimination of consciousness states

The initial investigation focused on determining whether any of the features or parameter ranges could effectively differentiate between different DoC states. We assessed whether features derived from experimental EEG signals, fitted NFT parameters, and features obtained from simulated EEG signals could act as reliable biomarkers for states of consciousness. Through statistical analysis using a non-parametric permutation test (10^4^ permutations) with false discovery rate multiple comparisons correction [48], many of these features and parameters demonstrated the capacity to differentiate between DoC and healthy individuals with statistical significance. However, none of the features or parameters were successful in differentiating between individual DoC states.

#### Features from experimental and simulated EEG

Figures 4a and 4b present features that exhibited significant differences between DoC and healthy states when extracted from both experimental and simulated EEG. The spectrum-related features Delta-power, Alpha-power, Beta-power, spectral slope, and spectral entropy prove effective discrimination in both the experimental and the simulated cases. This alignment was expected due to the CTM fitting process, which utilizes the spectrum. The entropy-related features also exhibit discriminative abilities between DoC and healthy states. Intriguingly, the discriminative power of the PE feature is statistically significant solely when calculated from experimental EEG data, whereas the LZc feature exhibits significance solely when computed from simulated EEG data.

**Figure 4.**
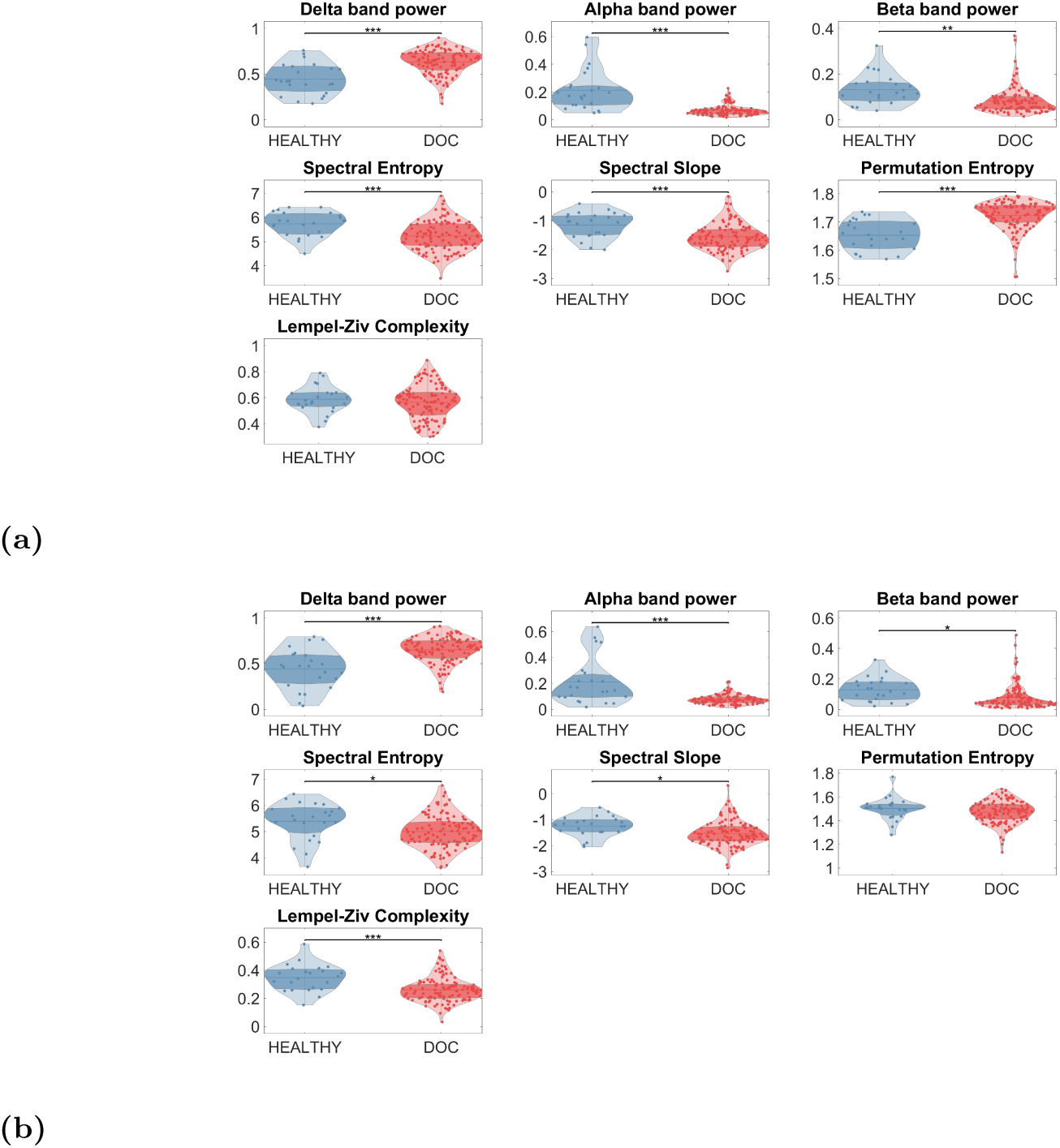
Differentiation of DoC states from a healthy state. The compared features were extracted from **(a)** experimental EEG and **(b)** CTM-simulated EEG.

#### NFT parameters

During this study, we observed that the repeated fitting of the CTM to experimental EEG data did not converge to a single set of parameters, in alignment with previous findings [26, 30]. Instead, it produced a broad range of parameter values with a complex distribution, while the goodness of fit (*χ*^2^) remained relatively stable (see Figure 5). Consequently, differentiating between states of consciousness could only be possible by using a sophisticated classifier. However, this approach is outside of the scope of the current work.

**Figure 5.**
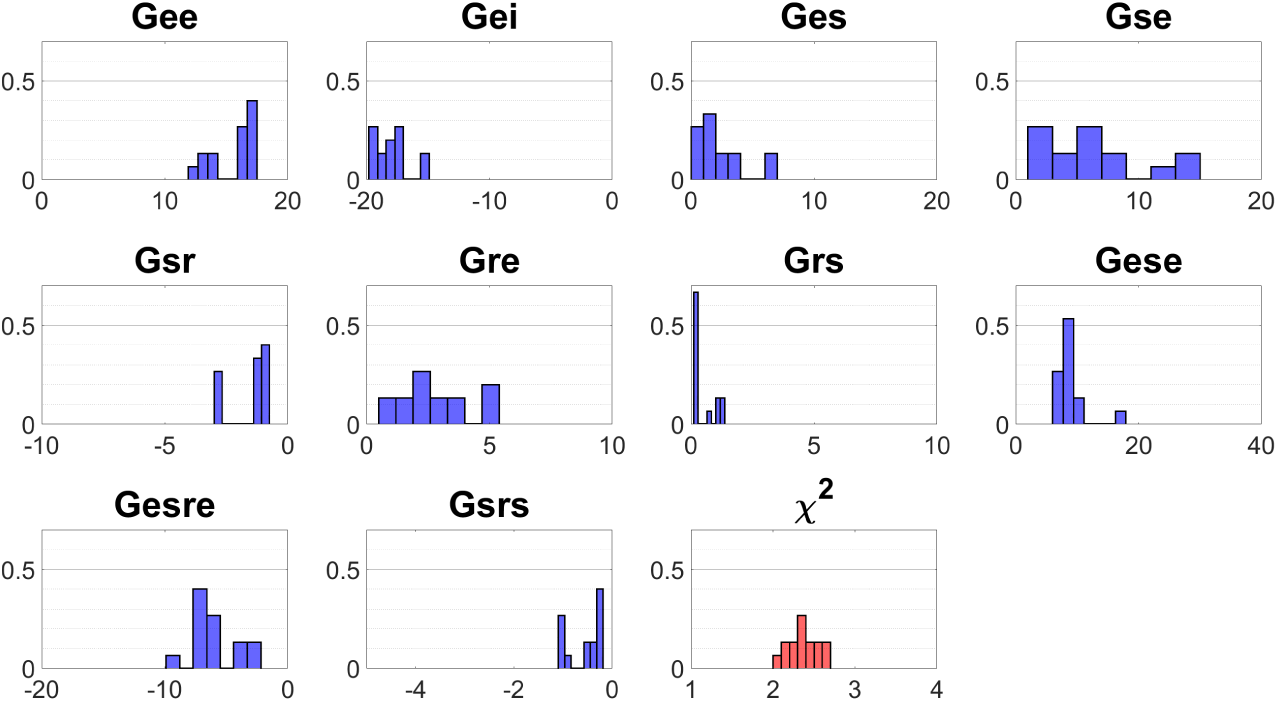
Distribution of NFT parameters obtained from 15 repeated fitting processes on a single MCS+ patient. Most parameters exhibited a wide distribution, covering a substantial portion of the typical range, and featuring several prominent peaks.

Nonetheless, we aimed to identify typical parameter value ranges for different DoC states. To achieve this, we repeated the fitting process 15 times for all subjects. Figure 6 illustrates NFT parameters with significantly distinct ranges for DoC and healthy states. (During the statistical analysis, we permuted all the 15 fits of a subject together.) Similar to the extracted features, the fitted NFT parameters could discriminate only between a healthy and DoC states, but not among different DoC states.

**Figure 6.**
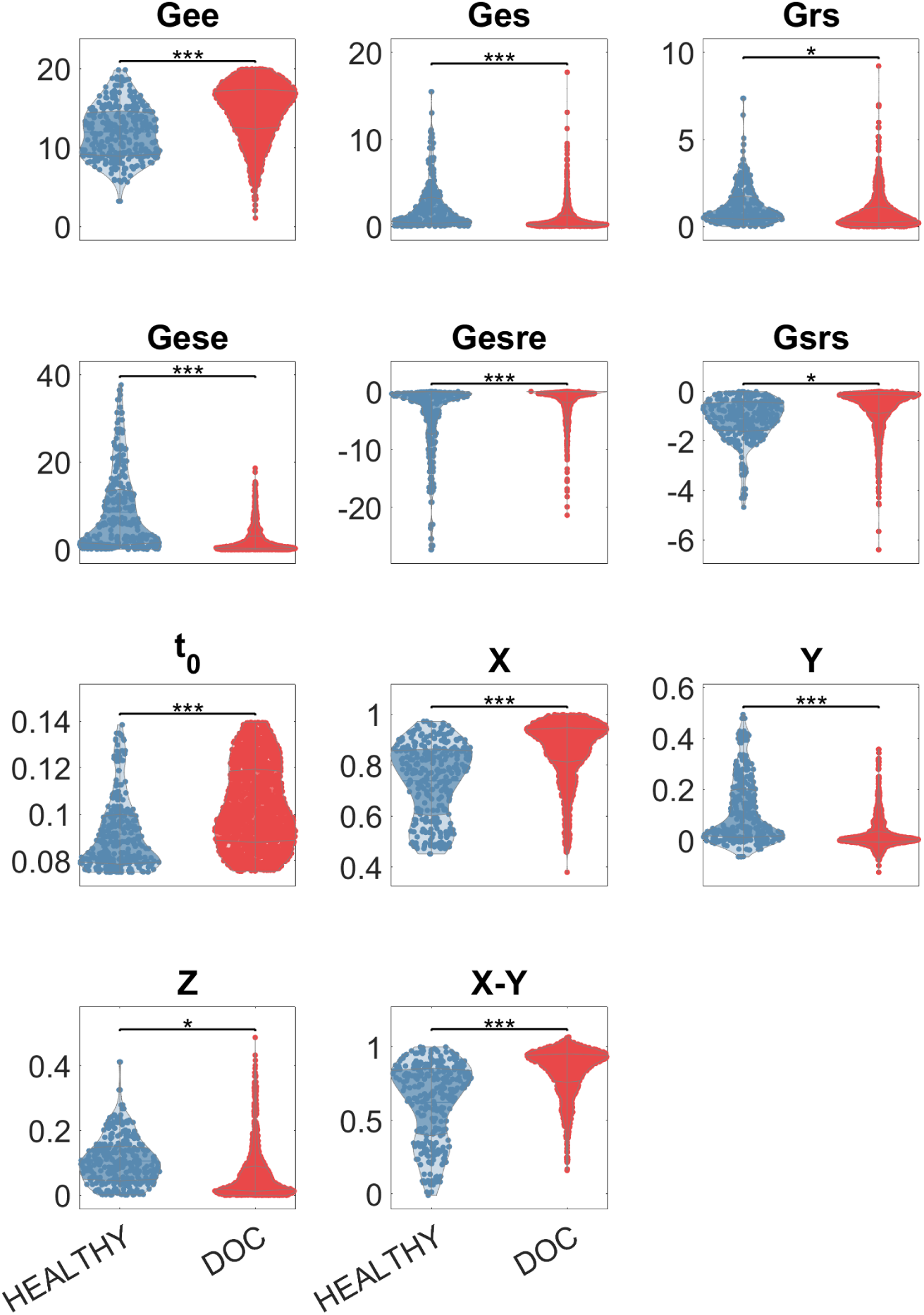
Differentiation of DoC states from a healthy state based on fitted NFT parameters. For each subject, 15 different sets of parameters were obtained during repeated fitting processes. Discriminatory capabilities primarily rely on combinations of parameters, such as *G_esre_* or *X,Y,Z*.

### Feature reproduction from simulated EEG

We explored correlations between features extracted from experimental EEG data and simulated EEG data. Notably, most of the spectrum-related features showed significant correlations. Specifically, Delta-power, Theta-power, Alpha-power, Beta-power, spectral slope, and spectral entropy features exhibited correlations with R*>*0.71 (*p <* 10*^−^*^23^), and their simulation-based values closely resembled the experiment-based results. In contrast, the Gamma-power feature did not display any significant correlations.

Among the entropy-related features, only LZc presented a substantial correlation of R=0.63 (*p <* 10*^−^*^16^), as depicted in Figure 7. While its simulation-based values were biased and exhibited shrinkage relative to the experiment-based ones, (i.e., the linear fit curve did not align with the *x* = *y* line), a simple linear transformation could rectify this misalignment.

**Figure 7.**
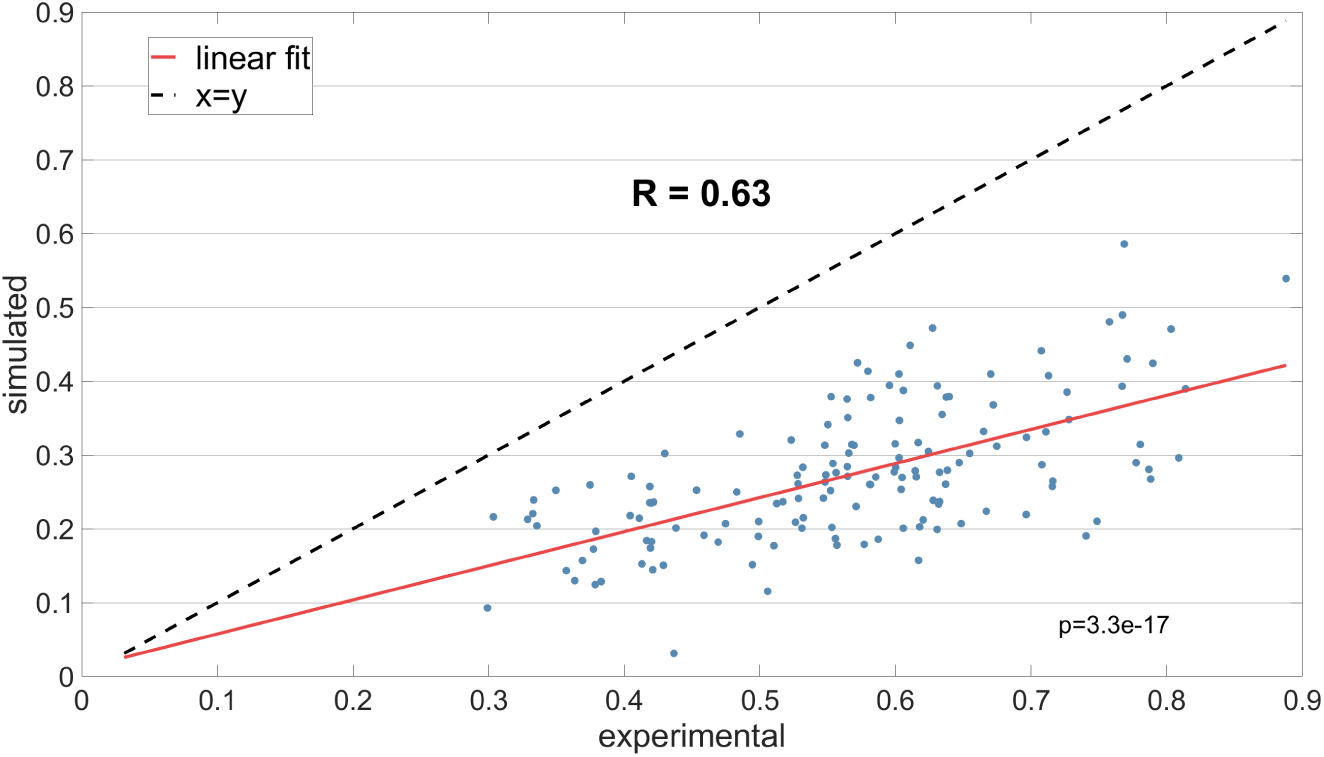
Correlation between LZc derived from experimental EEG data and LZc obtained from EEG data generated from a fitted CTM. The linear fit line illustrates the relationship between the two sets of data.

### Parameter-feature correlations analysis

We discovered significant correlations (p*<*0.001) between fitted parameters and features extracted from EEG generated by a fitted CTM. The Pearson correlation coefficients **R** for these relationships are provided in Table 2. Notably, NFT parameters that presented many correlations with the features tend to be stability-related (*X,Y,Z*, etc.).

**Table 2.**
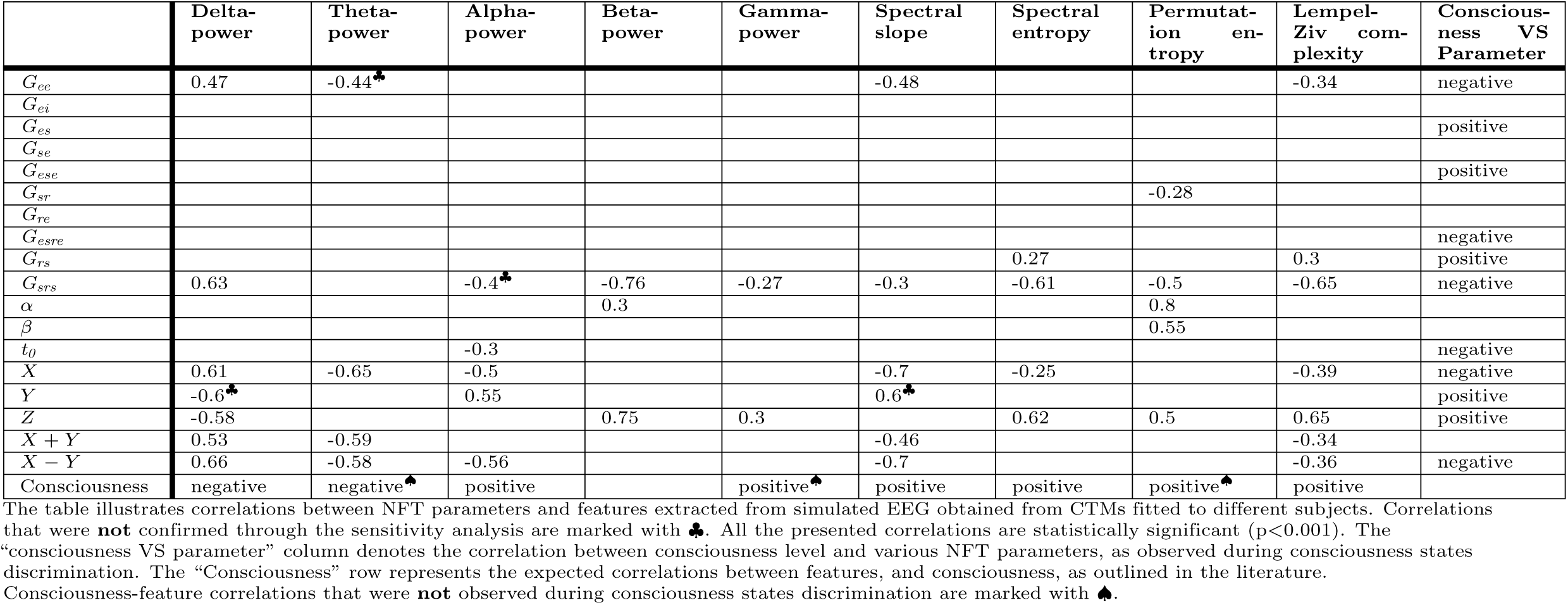
Parameter-feature correlations.

The majority of these correlations were validated through sensitivity analysis. Both of the representative subjects showed similar trends, with correlation coefficients maintaining the same sign and high statistical significance (p*<*0.05). Some of the correlations were not confirmed (indicated by ♣ in Table 2), but they were not contradicted either. In these cases, correlations were either insignificant or did not exhibit a clear monotonic trend, but neither did the healthy subject nor the MCS-patient display monotonic trends with opposite signs to the ones observed in the all-subjects correlation analysis. For instance, no significant correlation was observed between the excitatory cortical gain *G_ee_* and the Theta-power feature during the sensitivity analysis of a healthy subject. Interestingly, the trends comparison based on the sensitivity analysis revealed strong correlations that were not evident in the initial analysis with all subjects, such as those between *G_ee_*and Beta-power (see Figure 8). The reasons for these discrepancies are explored in the Discussion section.

**Figure 8.**
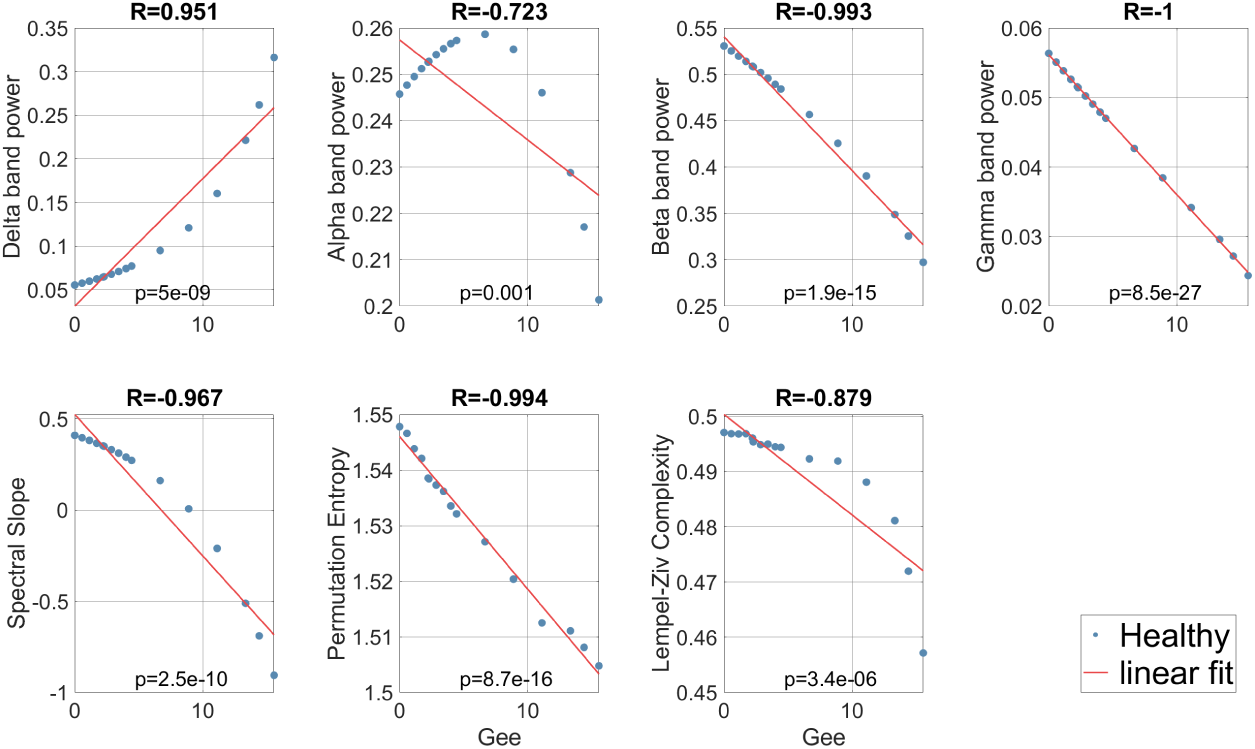
*G_ee_* parameter sensitivity of features extracted from simulated EEG. The CTM was fitted to a healthy subject. Only correlations with p*<*0.05 are presented.

Table 2 also includes the expected correlation trends of NFT parameters. In the section related to differentiating consciousness states, we showed that *G_es_*, *G_ese_*, *G_rs_*, *Y*, and *Z* are positively correlated with the level of consciousness, whereas *G_ee_*, *G_esre_*, *G_srs_*, *t_0_*, *X* and *X* − *Y* are negatively correlated with it.

Furthermore, in the table, we included the expected correlations of various features with the levels of consciousness, as reported in the existing literature. Previous research has indicated a positive correlation between the level of consciousness and the features: Alpha-power, Gamma-power [4], spectral slope [6, 49], spectral entropy [2, 9], PE, and LZc [15]. Conversely, a negative correlation has been observed between consciousness level and the features: Delta-power and Theta-power [4]. However, almost half of these features (marked with ♠) did not exhibit significant discrimination abilities between consciousness states, as demonstrated above. Furthermore, the experiment-based permutation entropy of DoC states was significantly higher than that of healthy states, contradicting previous findings [15]. This discrepancy discredits the correlation between simulation-based PE and consciousness level.

All of the identified correlations align with the expected correlation trends stemming from theoretical foundations, prior studies, and the results of consciousness state discrimination. For instance, we observed a positive correlation between spectral entropy and *Z*, which matches the positive correlation between consciousness level and both of them. However, the spectral entropy correlation with *X* was negative, which corresponds to a positive correlation between consciousness level and spectral entropy on one hand, and a negative correlation between consciousness level and *X* on the other hand. In general, we expected that parameter-feature correlations would exhibit positivity when both consciousness-feature and consciousness-parameter correlations are either positive or negative, and exhibit negativity when one of these correlations is positive while the other is negative. It is noteworthy that correlations between consciousness and features that were not confirmed by consciousness states discrimination (marked with ♠) suggest that the dataset did not reflect the anticipated relations from existing literature and were thus not utilized to test for mismatching.

## Discussion

In this work, we have delved into the application of NFT modeling in the study of neural correlates of consciousness. Despite its inherent simplicity as a brain model, NFT has shown promising potential for representing consciousness states. The model’s ability to capture consciousness states was evident through fitted NFT parameters and features based on simulated EEG, which exhibited discrimination capabilities and significant correlations. The NFT parameters also offered physiological interpretations for features based on simulated EEG and the states of consciousness. These various feature and parameter correlations underscore the suitability of NFT for advancing research in the field of consciousness, serving both as a tool for hypothesis generation and for conducting *in-silico* experiments.

### Feature-based discrimination of consciousness states

Many of our results align with previous findings. It was established in prior research that Alpha-power, spectral slope, and spectral entropy tend to have higher values in states of high consciousness and lower values in states of low consciousness, while the opposite is observed for Delta-power [2–4, 6, 9, 49]. In our study, we observed these behavior in both the features extracted from experimental EEG data and those derived from simulated EEG data. This alignment was expected, given that we employed the CTM’s power spectrum for model fitting (see Figure 3).

The LZc feature results were consistent with prior research, showing higher LZc values associated with higher consciousness levels and lower values with lower consciousness levels [15]. Interestingly, only the simulated-EEG-based LZc exhibited statistical significance (*p <* 0.001) in discriminating consciousness states. However, we showed that experimental-EEG-based LZc has a correlation of R=0.63 with simulated-EEG-based LZc. A potential explanation for this disagreement could be that experimental data may contain noise and biases that impact the LZc feature, which are filtered out during the fitting process. This implies that the fitted CTM might emphasize LZc as a potential biomarker for its inherent consciousness state.

Intriguingly, we observed a divergence from existing literature concerning the PE feature, with lower values observed in healthy subjects and higher values in DoC patients. This contradicts previous findings, which typically report the opposite trend [15]. Unfortunately, we were unable to provide a definitive explanation for this discrepancy. Further investigation is warranted to elucidate the underlying reasons for this unexpected finding.

It is important to note that the ability to distinguish between healthy and DoC states, while not being able to differentiate among DoC sub-states, aligns with expectations for these features. The classification of DoC sub-states based on EEG features remains a challenging task, with limited success achieved so far, typically in conjunction with a preceding transcranial magnetic stimulation [13]. Therefore, the capability to differentiate between healthy and DoC states based on resting-state EEG data, whether experimental or simulated, is a noteworthy achievement.

### Parameter-based discrimination of consciousness states

We successfully replicated the main findings of similar research previously conducted by Assadzadeh et al. (2023). Assadzadeh used the same DoC database, fitted CTMs to the subjects, and applied a linear discrimination analysis on NFT parameters to classify various DoC and healthy states. By combining *X* − *Y*, *X* + *Y*, and 1*/α* + 1*/β*, they achieved a classification accuracy of 93% for distinguishing between DoC and healthy states, but not among different DoC states [31]. While classifying among consciousness states was not within the scope of the current study, a combination of various features and NFT parameters could potentially yield promising results in this regard.

Instead, our focus was on understanding how each of the NFT parameters reflects the state of consciousness, and our findings were consistent with those presented by Assadzadeh et al. We discovered, akin to their results, that the excitatory cortical gain *G_ee_*, corticothalamic loop gains *G_ese_* and *G_esre_*, thalamothalamic loop gain *G_srs_*, and the stability parameters *X,Y,Z*, and *X* − *Y*, all exhibited significantly divergent ranges of values (p*<*0.05) between DoC and healthy states. Additionally, we identified significant discriminatory capabilities for consciousness states through the *G_es_*, *G_rs_*, and *t_0_* parameters. This suggests that each of these parameters contributes to the representation of the level of consciousness within the CTM, providing distinct physiological insights into varying consciousness states.

It is essential to bear in mind that while we identified typical parameter ranges for DoC and healthy subjects, we should not depend on a single parameter value for determining the state of consciousness. As demonstrated, CTM parameters can converge to different values for the same subject. We speculate that these different values, when combined with other parameters, form a manifold that represents a state of consciousness. It is possible that parameter combinations such as *X* − *Y* partially describe this manifold, providing them with the capability to differentiate between states of consciousness.

### Features reproduction

We aimed to investigate whether a CTM fitted to a particular subject can produce EEG features with values that closely match those obtained from experimental EEG. Our analysis identified strong correlations between all the spectrum-related features, except Gamma-power, extracted from both experimental EEG and simulated EEG data. Additionally, a notable correlation was observed between the LZc feature derived from both types of data. The strong correlation and the close match between the experiment-based and simulation-based spectrum-related feature values was anticipated, given that the CTM was fitted based on the power spectrum. The lack of correlation for Gamma-power might be explained by the fitting objective function, which assigns more weight to lower frequency power than to higher frequency power [30]. The impact of this weighting is evident in Figure 3, where the frequencies in the Gamma band of the analytical and simulated spectra do not follow the experimental spectrum.

The correlation discovered for LZc is quite interesting and meaningful. It suggests that, in EEG-based research, simulated EEG can be used as a viable alternative to experimental EEG when employing LZc as a measure. Furthermore, if a CTM is utilized for *in-silico* experiments, LZc can act as a valuable biomarker for consciousness states, paralleling its extensive use in prior studies on NCC. This finding also indicates that the fitted CTM has the capability to control the complexity of the generated signal using its parameters. We present these parameters in Table 2, offering a physiological interpretation for LZc.

### Parameter-feature correlations

In this part, our aim was to investigate how each fitted parameter influences the features computed from simulated EEG. As mentioned earlier, most of the features exhibit correlations with NFT parameters, and these correlations offer valuable insights into the physiological explanations for each feature. For instance, we found that PE is moderately affected by *G_sr_*, *G_srs_*, *NFT α*, *NFT β*, and *NFT Z* which represent the thalamic reticular-relay nuclei connection gain strength, thalamothalamic loop gain strength, synaptodendritic decay and rise rates, and intrathalamic loop strength, respectively. The positive correlations between PE and *NFT α* and *β* make intuitive sense: higher synaptodendritic response rates lead to faster alterations in the output signal, resulting in a more complex and informative signal, which translates to higher entropy.

A notable point from Table 2 is the dominant role of stability-related parameters *X, Y, Z*, and their combinations in driving the majority of correlations observed. This aligns with prior research suggesting that *X,Y*, and *Z* qualitatively encapsulate a significant portion of CTM’s dynamics [28, 30]. This observation supports our hypothesis that a single parameter may not fully represent a specific model behavior; instead, a set of parameters, potentially forming a complex manifold, might be required. However, this does not diminish the importance of correlations involving other individual parameters, which also play a critical role in elucidating various features, as previously discussed.

We compared our results to those from parameter sensitivity analyses in previous studies. Robinson et al. (2001) explored how different parameters affect the analytical power spectrum of a simplified CTM (excluding the thalamic reticular nucleus) fitted to a healthy, awake, eyes-closed subject [32]. Subsequently, Rowe et al. (2004) performed a similar analysis on the complete CTM, replicating Robinson’s findings and exploring additional parameters [36]. Most of the correlations we identified between spectrum-related features and NFT parameters were consistent with these studies (except for spectral entropy, which was not analyzed by Robinson and Rowe). However, some correlations reported in previous studies are not evident in our results. These include correlations of *G_ee_* with Alpha-power, *t_0_* with Delta and Theta-power, *NFT α* with Gamma-power, and all correlations involving *G_ei_*, *G_ese_*, *G_esre_*, and *NFT β*. Additionally, we identified correlations (albeit low) of *G_srs_* with Gamma-power and spectral slope that were not reported previously. These discrepancies might stem from the significantly different experimental conditions between our study and previous ones, as well as from our reliance on simulated data of multiple subjects versus the analytical spectrum of a single subject used in the earlier research.

In the sensitivity analysis, we explored the parameter space around a fitted parameter set. This parameter corresponds to a fixed point on a manifold, representing all possible parameter combinations that produce a fitted spectrum. Such fixed points on the manifold may exhibit varying degrees of sensitivity to individual parameters. By selecting a different point on the manifold, or simply by exploring the fitted parameters of another subject, we may encounter heightened sensitivity to certain parameters, while observing reduced sensitivity to others. The overall trend encompasses various fixed points for all of the different subjects, yet not all of these points demonstrate sensitivity to the same parameters.

This perspective can also offer an explanation for correlations that appeared during the sensitivity analysis, but were absent in Table 2. For example, the correlation between *G_ee_* and Beta-power, as seen in Figure 8, reflects the behavior of parameters around a fitted parameter set of a particular healthy subject. However, this behavior is not universal across all subjects, indicating that the correlation is subject-specific and not generalizable. Consequently, the examination of trends through the sensitivity analysis should ideally be extended to encompass all subjects within the dataset to validate the overall parameter-feature correlations, as previously emphasized.

The alignment of the discovered correlations with the anticipated trends from NCC and the consciousness states’ discrimination results underscores the NFT model’s potency to represent different levels of consciousness. The reliable triadic relationship between parameter values, consciousness levels, and feature values means that both a fitted NFT parameter and a simulated-EEG-based feature can effectively represent the consciousness state, with changes in one being reflected in the other.

### Consciousness state representation in the NFT model

The CTM effectively reflects the state of consciousness through features derived from simulated EEG data. These features are also influenced by NFT parameters, which exhibit distinct value ranges corresponding to different states of consciousness. This triadic relationship offers a strong validation that the model faithfully mirrors the specific state of consciousness of the subject to which it is fitted.

A particularly interesting example is the LZc feature, which effectively discriminates between subjects in different states of consciousness. It correlates with *G_srs_*and stability-related NFT parameters, all of which demonstrate the capability to distinguish between different consciousness states as well. Moreover, a robust correlation exists between simulated-EEG-based LZc and experimental-EEG-based LZc. Thus, LZc extracted from simulated data emerges as a reliable biomarker for assessing the consciousness level within the CTM, akin to its utility in previous NCC-related studies.

The compelling evidence presented in this study lends credibility to the use of NFT as a valuable tool in consciousness research for applications that include model-based interpretations, hypothesis generation, and *in-silico* experiments. For example, consider a scenario where researchers aim to investigate the effects of a drug that suppresses intrathalamic feedback mechanisms on the level of consciousness. A CTM could be fitted to a healthy subject, and *G_srs_* could be manipulated to generate an EEG time series. Then LZc for various *G_srs_*values could be calculated, and the consciousness level could be determined accordingly. In another scenario, if a subject has been administered propofol and exhibits an observable increase in Delta-power, the fitted CTM may suggest that the subject’s condition can be explained by high corticocortical loop strength *X*. However, Delta-power is also influenced by *G_srs_*. Therefore, researchers may hypothesize that administering the studied drug to the propofol-anesthetized subject will reverse the observed effects and potentially wake her/him up.

Overall, the utilization of a reliable computational brain model in consciousness research offers numerous advantages, whether seeking model-based predictions or investigating the neural mechanisms underlying a phenomenon. When it comes to experiments involving live subjects, the model helps to avoid potential health risks and negative implications. Furthermore, for other experiment types and theoretical work, computational modeling can significantly save time, effort, and expenses. The prospects of integrating NFT modeling into consciousness research are promising. We anticipate that future research endeavors will increasingly harness NFT to gain deeper insights into the intricate nature of consciousness and contribute to advances in the field of NCC. Yet, a more immediate research direction should include a similar exploration of data from other consciousness-related states, such as sleep, anesthesia, and psychedelics, to inspect how the CTM represents them and to enhance our understanding of the brain’s functioning across these states of consciousness.

## Summary

In this study, we explored the application of NFT modeling in investigating and understanding NCC. Despite its inherent simplicity, NFT has shown considerable promise in representing consciousness states in the brain.

Our primary findings indicate that CTM is capable of reflecting consciousness states, both in terms of fitted NFT parameters and features derived from simulated EEG. These features have exhibited discrimination capabilities and have shown significant correlations with model parameters, affirming CTM’s potential as a model for understanding and representing consciousness.

Our results suggest that NFT can be a valuable tool in consciousness research, supporting model-based interpretations, hypothesis generation, and *in-silico* experiments. This study establishes a strong foundation for further exploration of NFT, offering new insights into consciousness and the brain’s dynamics. By deepening our understanding of these relationships, this research could inspire innovative approaches in neuroscience, cognitive science, and consciousness studies.

## Statements

### Compliance with ethical standards

The study was approved by the ethical committee at the Medicine Faculty of the University of Liege, Belgium, in accordance with the Declaration of Helsinki. All healthy subjects gave written informed consent, while for non-communicating brain-injured patients, the informed consent was obtained from a legal surrogate.

### Data availability

The data underlying this article are available on Zenodo with the identifier 10.5281/zenodo.13919013.

## Acknowledgments

We would like to thank Dr. Eli J. Müller from the Brain and Mind Centre at the University of Sydney for providing invaluable insights into the NFT framework software.

This work was supported by the University and University Hospital of Liège, the Belgian National Funds for Scientific Research (FRS-FNRS), the FNRS PDR project (T.0134.21), the ERA-Net FLAG-ERA JTC2021 project ModelDXConsciousness (the Human Brain Project Partnering Project), and the FNRS MIS project (F.4521.23), O.G. is a research associate at FRS-FNRS.

The wording revision process of this article included the use of ChatGPT3.5 by OpenAI.

